# Evolutionary conservation of the grape sex-determining region in angiosperms and emergence of dioecy in *Vitaceae*

**DOI:** 10.1101/2024.11.10.622717

**Authors:** Mélanie Massonnet, Noé Cochetel, Valentina Ricciardi, Andrea Minio, Rosa Figueroa-Balderas, Jason Londo, Dario Cantu

**Affiliations:** Department of Viticulture and Enology, University of California Davis, Davis, CA, 956161, USA; Dipartimento di Scienze Agrarie e Ambientali, Università degli Studi di Milano, Milan, Italy; School of Integrative Plant Science, Cornell University, Ithaca, NY, 14850, USA; Genome Center, University of California Davis, Davis, CA, 95616, USA

**Keywords:** Flower sex determination, dioecy, recombination suppression, angiosperms, comparative genomics

## Abstract

Dioecy, defined by the presence of either female or male flowers on separate individual plants, is rare in angiosperms. However, it has independently evolved multiple times from hermaphroditic ancestors. In *Vitis* spp., flower sex is determined by a ∼200-kbp region with five genes involved in flower development and sexual identity. This study traces the evolutionary history of the *Vitis* sex-determining region (SDR) to assess whether it evolved in a conserved flowering-related locus. We analyzed the conservation of the *Vitis* SDR across 42 plant genomes and found collinearity in all 39 angiosperms, but not in non-flowering plants. The number of collinear genes and their rank-normalized gene conservation score were higher in the *Vitis* SDR compared to the rest of the genome, indicating its importance for flowering. We further explored SDR conservation within the *Vitaceae* family by long-read sequencing and haplotype phasing of eight species, including the outgroup *Leea coccinea*. Similarly, the *Vitis* SDR was highly conserved across the *Vitaceae* genomes, although variations in region size, primarily attributed to differences in repetitive elements, were observed. Interestingly, no recombination suppression was found in the dioecious *Tetrastigma*, suggesting a different sex determination mechanism. In *Muscadinia rotundifolia*, linkage disequilibrium analysis, haplotype comparison, and gene expression profiling showed that muscadine grapes have similar SDR boundaries and candidate sex-determining genes as *Vitis*. These results suggest that dioecy emerged in a common ancestor of the two grape genera at a locus associated with flower morphology and fertility, highly conserved in flowering plants.

## Introduction

Dioecy, the presence of distinct male and female individuals within a species, is rare, occurring in approximately only 6% of flowering plants (Renner, 2014). Dioecy has evolved independently multiple times from hermaphroditic ancestors in angiosperms, with estimates suggesting between 900 and 5,000 transitions across 175 plant families (Renner, 2014). Interestingly, dioecy often reverts back to hermaphroditism, underscoring the remarkable plasticity and adaptability of plant reproductive strategies (Renner, 2014; Goldberg et al., 2017).

In *Vitis* spp., or bunch grapes, all ∼80 species are dioecious, except for the domesticated grapevine (*V. vinifera* ssp. *vinifera*), which produces hermaphroditic flowers (Moore, 1991). In *Vitis* spp., flower sex is determined by a sex-determining region (SDR) located on chromosome 2 and spanning approximately 200 kbp (Massonnet et al., 2020; Zou et al., 2021). The *Vitis* SDR exhibits a high linkage disequilibrium with boundaries strictly conserved across the genus (Massonnet et al., 2020; Zou et al., 2021). Divergence time estimates suggest that dioecy in *Vitis* originated at least ∼20 million years ago (Mya) (Zou et al., 2021). The *Vitis* SDR is composed of twelve to fourteen genes (Zou et al., 2021), with five genes involved in floral organ development and identity in plants (**Table 1**). These include the two candidate sex-determining genes *VviYABBY3* and *VviINP1* (Massonnet et al., 2020; Zou et al., 2021), and *VviPLATZ1*, which has been shown to influence stamen morphology (Iocco-Corena et al., 2021). The multiple closely linked flowering-related genes within the SDR raise questions about whether this region is unique to the *Vitis* genus or if it is also present in the broader *Vitaceae* family, or even in more distantly related plants.

**Table 1:** Homologs of the *Vitis* SDR genes have a role in flower development and sex fertility.

| Functional description | Plant species | Gene name | Mutant | Phenotype | Reference |
| --- | --- | --- | --- | --- | --- |
| YABBY transcription factor | *Arabidopsis thaliana* | *AtYABBY1*,<br>*AtFIL*,<br>*AtAFO* | Knockout *fil* | Altered floral organ number, position, identity and development during flower development. Partial male sterility. | (Sawa et al., 1999) |
|  | *Oryza sativa* | *OsYABBY4* | CAM35S-*OsYABBY4* | Increased number of stamens, pistils and stigmas. | (Yang et al., 2016) |
| Vacuolar invertase | *Arabidopsis thaliana* | *AtBETAFRUCT4* | Knockouts *Atvi2-1*,<br>*Atvi2-2* | Reduced pollen germination. | (Seitz et al., 2023) |
| SKU5 | *Arabidopsis thaliana* | *AtSKU5* | Knockouts *sks1*,*sks2*,*sks3*,*sku5* | Smaller flowers than wild type. Reduced seed production due to maternal-dependent ovule abortions. | (Zhou, 2019) |
| INAPERTURATE POLLEN1 | *Arabidopsis thaliana* | *AtINP1* | Knockout *atinp1* | Complete lack of pollen aperture. | (Dobritsa et al., 2011) |
|  | *Oryza sativa* | *OsINP1* | Knockout *osinp1* | Complete lack of pollen aperture. Sterile pollen. | (Zhang et al., 2020) |
|  | *Zea mays* | *ZmINP1* | Knockout *zminp1* | Complete lack of pollen aperture. Sterile pollen. | (Li et al., 2018) |
|  | *Eschscholzia californica* | *EcINP1* | Silencing pTRV2:*EcINP1* | Abnormal apertures. | (Mazuecos-Aguilera et al., 2021) |
| PLATZ transcription factor | *Vitis vinifera* | *VviPLATZ1* | Knockout *vviplatz1* | Reflex filaments. | (Iocco-Corena et al., 2021) |

The *Vitis* genus belongs to the Viteae clade within the *Vitaceae* family (**Fig. 1)** (Sun et al., 2016; Ma et al., 2021). The *Vitaceae* family is divided into five major clades: (i) Ampelopsideae, (ii) Parthenocisseae, (iii) Viteae, (iv) Cayratieae, and (v) Cisseae (Wen et al., 2018). The family comprises 16 genera and ∼950 species distributed across tropical and temperate regions, characterized by their distinctive leaf-opposed tendrils (Wen et al., 2018). *Vitaceae* exhibit significant morphological diversity, particularly in vegetative structures, seed characteristics, inflorescence types, and floral features (Gerrath et al., 2017). There is also considerable variation in chromosome numbers, ploidy levels, and genome sizes both among and within *Vitaceae* clades (Karkamkar et al., 2011; Chu et al., 2018). For example, *Vitis* ssp. are diploid with 19 chromosomes (2n = 38), individuals of the second grape genus (*Muscadinia* spp.) are diploid with 40 chromosomes (2n = 40), and some *Cissus* species exhibit tetraploidy or hexaploidy (Chu et al., 2018). While most *Vitaceae* species are hermaphroditic, other mating systems, including dioecy, also occur. Dioecy is found within the Viteae clade, which includes the grape genera *Vitis* and *Muscadinia*, as well as the *Tetrastigma* genus of the Cayratieae clade (Wen et al., 2018).

**Fig. 1:**
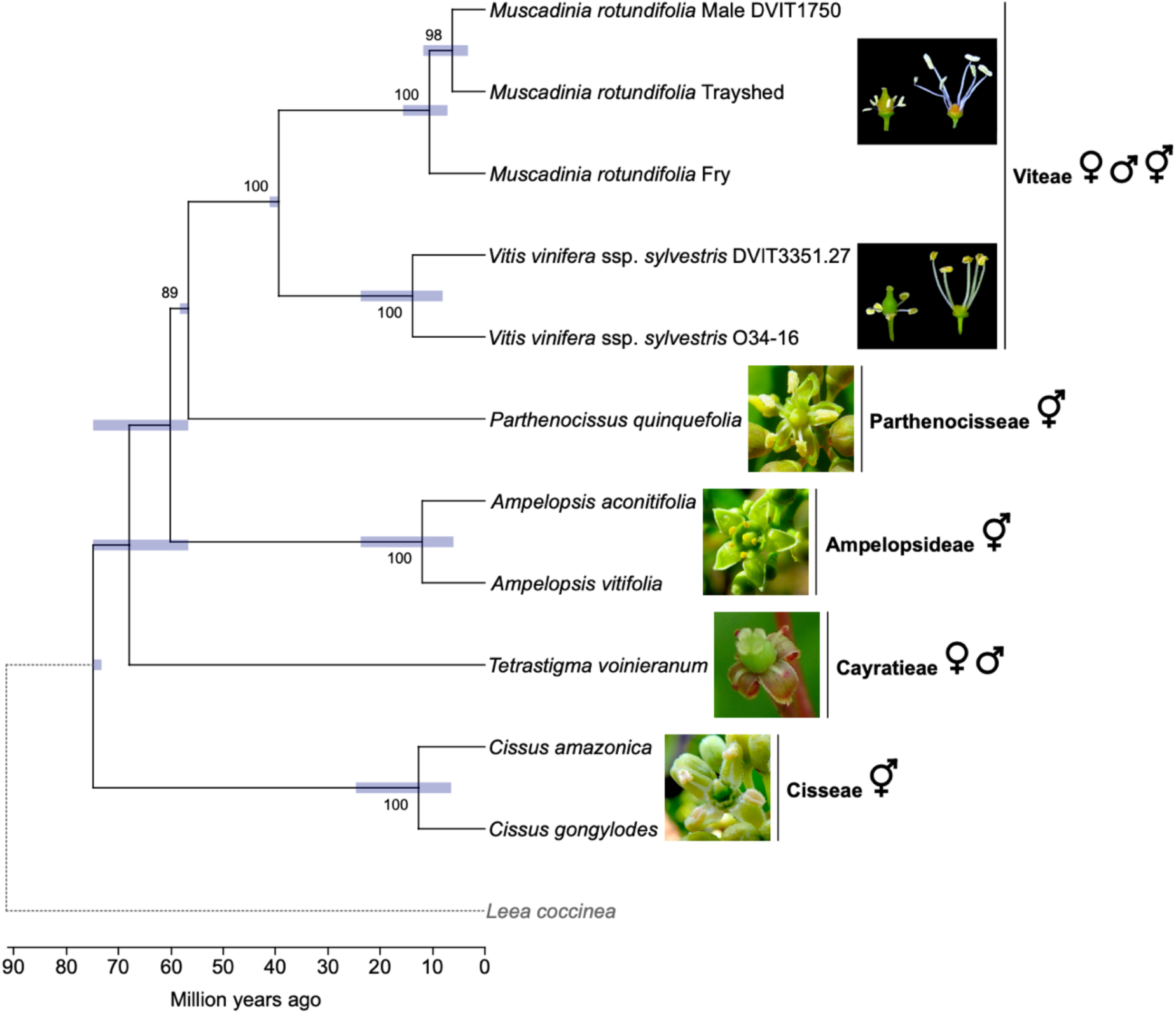
Phylogeny of the *Vitaceae* and morphology of their flowers. Phylogenetic tree predicted from single-copy orthologues in angiosperms (Johnson et al., 2019). The branches represent divergence times in million years. Bars around each node represent 95% confidence intervals. *L. coccinea* was used as an outgroup to the *Vitaceae* family. Bootstrap values greater than 80 are indicated. The symbols ♀, ♂ and ⚥ represent the female, male, and hermaphrodite flower sex type present in each clade, respectively. Picture of a female flower from *Tetrastigma* was modified from Habib *et al*. (Habib et al., 2017).

*Vitis* and its sister genus *Muscadinia* diverged approximately 18-47 Mya (Wan et al., 2013; Liu et al., 2016; Ma et al., 2018) and each consist of dioecious species (Biasi and Conner, 2016). A high-density linkage map has located the SDR in *M. rotundifolia* on the same chromosomal region as in *Vitis* spp. (Lewter et al., 2019), suggesting that both genera have evolved from a common dioecious ancestor. However, the precise boundaries of the muscadine SDR have yet to be defined, making it unclear whether the same genes regulate flower sex in both grape genera. A previous study identified the same candidate male-sterility mutation in the female associated (F) haplotype of the male *M. rotundifolia* cv. Trayshed as in *Vitis* species (Massonnet et al., 2020). However, for the candidate female-suppressing gene in *Vitis* spp., *VviYABBY3*, Trayshed lacked the two non-synonymous SNPs specific to the male-associated (M) allele in *Vitis*, and the gene expression of *VviYABBY3* in muscadine flowers has yet to be investigated.

In this study, we explored the evolutionary history of the *Vitis* SDR over approximately 200 million years within plants and the *Vitaceae* family. To achieve this, we examined gene collinearity and conservation of the *Vitis* SDR across 42 plant genomes, encompassing a broad range of angiosperms from monocots to eudicots. We also analyzed the conservation of the *Vitis* SDR among four other *Vitaceae* clades by sequencing and assembling genomes of six *Vitaceae* species and the outgroup *Leea coccinea*. Furthermore, we assessed whether the orthologous SDR region in *Vitis* could be associated with sex determination in the dioecious *Tetrastigma* species. Lastly, we investigated the muscadine grape SDR by defining its boundaries and identifying candidate sex-determining genes. This comprehensive analysis sheds light on the evolution and functional significance of the *Vitis* SDR across diverse plant taxa.

## Results

### The *Vitis* sex-determining region is highly conserved in flowering plants

To investigate the evolutionary history of the *Vitis* SDR in plants, we analyzed collinearity between the grapevine SDR and its orthologous regions across 42 plant genomes (**Supplementary Table 1**), comparing these results to overall genomic collinearity. For this analysis, we utilized the SDR from the haplotype 1 of *Vitis vinifera* ssp. *vinifera* cv. Cabernet Sauvignon (hereafter referred to as Cabernet Sauvignon). The dataset comprised 39 angiosperms (33 eudicots and six monocots) and three non-flowering species: the gymnosperm *Taxus chinensis* (Chinese yew), the tracheophyte *Selaginella moellendorffii* (spike moss), and the embryophyte *Physcomitrium patens* (spreading earthmoss).

In angiosperms, we identified an average of 1,863.3 ± 280.6 collinear windows, whereas non-flowering species showed significantly fewer windows (30.3 ± 21.2). Importantly, a collinear window corresponding to the SDR was detected in all angiosperms but was absent in the three non-flowering species (**Supplementary Fig. 1**). Among angiosperms, the number of genes within the collinear window of the *Vitis* SDR was significantly higher than those in collinear windows of the other 19 grape chromosomes (Kruskal-Wallis test; *P* = 2.0 × 10⁻²³; **Fig. 2a**). Additionally, the rank-normalized gene conservation score for the collinear genes within the *Vitis* SDR collinear window was significantly greater than that of other collinear genes (Kruskal-Wallis test; *P* = 7.1 × 10⁻¹³; **Fig. 2b**). Most genes in the SDR collinear window were conserved across angiosperm genomes, including both eudicots and monocots (**Fig. 2c**). Manual refinement of gene models identified 18 additional protein-coding genes and 14 pseudogenes within the SDR collinear window across 13 genomes (**Supplementary Fig. 1**; **Supplementary Table 2**). Notably, the two candidate sex-determining genes in grapes, *VviYABBY3* and *VviINP1*, were located within the same collinear window in 27 eudicots (81.8%) and the monocot *Phoenix dactylifera* (date palm; 16.7%; **Supplementary Fig. 1**). These findings collectively indicate that the genomic region corresponding to the *Vitis* SDR is highly conserved among angiosperms but absent in non-flowering species.

**Fig. 2:**
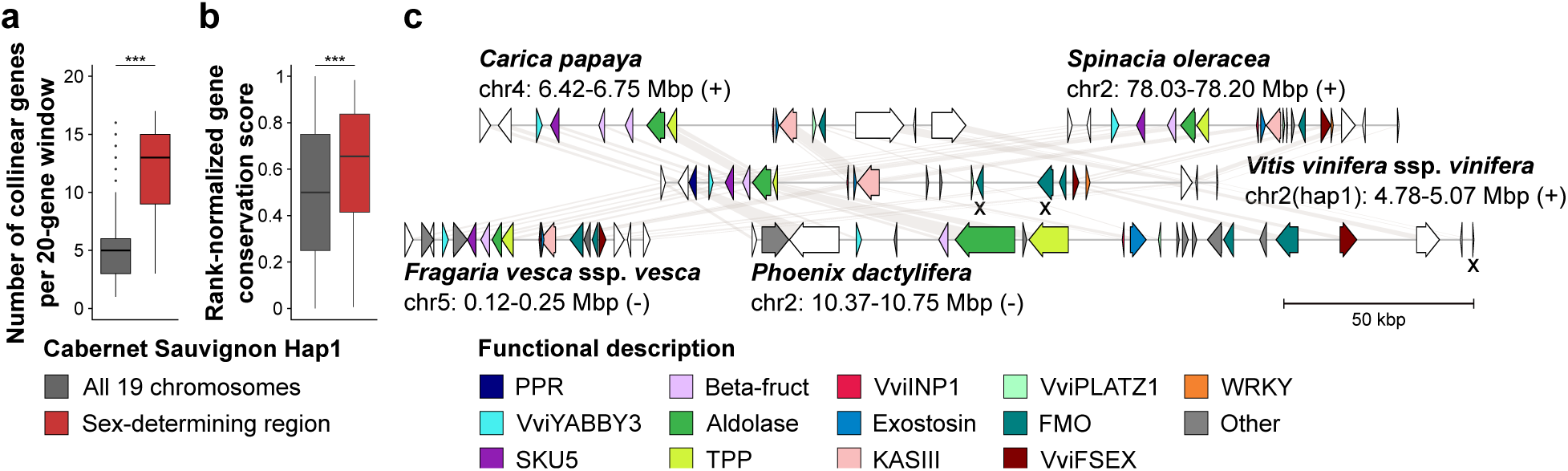
Conservation analysis of the *Vitis* sex-determining region (SDR) genes in 42 plant genomes. Number of collinear genes per 20-gene windows (*i.e*. 20-gene window with collinear genes) in the nineteen chromosomes of Cabernet Sauvignon Haplotype 1 (Hap1) and the SDR among the plant genomes. (**b**) Gene conservation score of the collinear genes in the Cabernet Sauvignon Haplotype 1 and the SDR among the angiosperm genomes. Panels (**a**) and (**b**) share the same color legend. ***, *P* value < 1 × 10^−11^. (**c**) Schematic representations of the gene content in the homologous regions of the *Vitis* SDR in four angiosperms. Each haplotype is annotated with arrows to depict annotated genes. White arrows indicate genes that are not part of the *Vitis* SDR. Genes affected by nonsense mutations are indicated with an X. The scale below the haplotypes denotes the length of the region.

### The *Vitis* sex-determining region is conserved across the *Vitaceae* family

To investigate whether the genomic region corresponding to the *Vitis* SDR is conserved across the *Vitaceae*, we sequenced, assembled, and phased the genomes of six *Vitaceae* accessions from four major clades: *Parthenocissus quinquefolia* (Parthenocisseae clade), *Ampelopsis aconitifolia* and *A. vitifolia* (Ampelopsideae clade), *C. amazonica* and *C. gongylodes* (Cisseae clade), and *T. voinieranum* (Cayratieae clade), as well as *L. coccinea* as outgroup (**Fig. 1**; **Supplementary Tables 3-5**). The genome assembly sizes for *A. vitifolia* and *C. amazonica* were 785.6 Mbp and 704.3 Mbp, respectively, while *A. aconitifolia*, *P. quinquefolia*, and *C. gongylodes* had larger genome sizes of 999.1 Mbp, 1,136.4 Mbp, and 1,270.6 Mbp, respectively (**Supplementary Table 5**). The diploid genome of *T. voinieranum* was the largest of the study, with a size of 4,419.2 Mbp. Regarding the outgroup *L. coccinea*, its diploid genome was 1,104.9 Mbp long.

By aligning the SDR genes from Cabernet Sauvignon, we identified 21 regions homologous to the *Vitis* SDR: two in the outgroup *L. coccinea*, two in *P. quinquefolia*, two in *A. aconitifolia*, one in *A. vitifolia*, four in *C. amazonica*, eight in *C. gongylodes*, two in *T. voinieranum* (**Fig. 3**). The number of haplotypes found in *C. amazonica* and *C. gongylodes* suggest that these two species are tetraploid and octoploid, respectively. In *A. vitifolia*, only one *Vitis* SDR-homologous region (VSR) was identified. Alignment of short DNA sequencing reads from this accession revealed 160 heterozygous SNPs within the region (i.e., 0.05 SNPs per kbp), indicating low heterozygosity, which may explain why only one haplotype was generated by FALCON-Unzip.

**Fig. 3:**
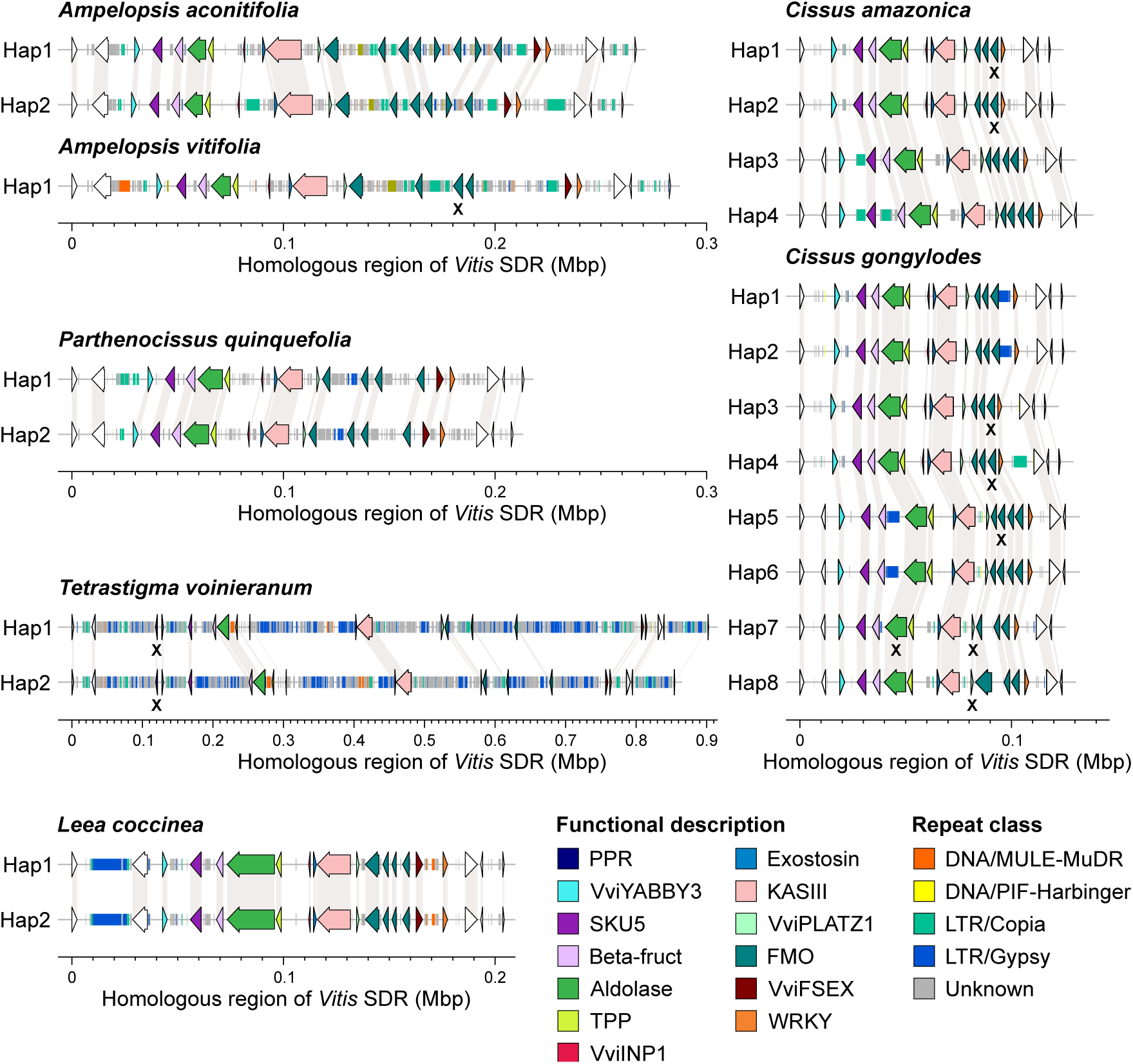
Schematic representations of the gene and intergenic repeat contents in the homologous regions of the *Vitis* sex-determining region in Vitales. Each haplotype is annotated with arrows to depict annotated genes. White arrows indicate genes that are not part of the *Vitis* sex-determining region. Genes affected by nonsense mutations are indicated with an X.

The gene content across the identified haplotypes was largely conserved (**Fig. 3**). We could observe some differences between clades and even species within clades. For example, in *A. aconitifolia*, we identified seven and five genes encoding flavin-containing monooxygenases (FMOs) in the first and second haplotypes, respectively, while *A. vitifolia* had four FMO genes in its single haplotype. The number of FMO genes also varied between the two *Cissus* species, and the gene encoding the hypothetical protein *VviFSEX* was absent in all *Cissus* haplotypes. Additionally, the *VviINP1* gene was present in half of the haplotypes in both *C. amazonica* and *C. gongylodes*. We further investigated whether the deleterious 8-bp deletion in the candidate sex-determining gene *VviINP1* was present in other *Vitaceae*. Alignment of DNA-seq short reads from 159 *Vitaceae* accessions, representing 88 species, revealed that this mutation was only found in the *Vitis* genus and *Muscadinia rotundifolia* (**Supplementary Table 6**).

Despite the high conservation of the gene content, the size of the VSRs varied significantly between clades, species, and haplotypes. The region from the NAC transcription factor gene to the gene homologous to *Arabidopsis thaliana PURPLE ACID PHOSPHATASE 2* (PAP2) was the shortest in the two *Cissus* species (122.6 ± 3.9 kbp), followed by *P. quinquefolia* (213.4 and 208.3 kbp), the two *Ampelopsis* species (270.0 ± 11.5 kbp), and *T. voinieranum* (902.6 and 854.6 kbp). In the outgroup *L. coccinea*, both haplotypes were 204.3 kbp, and the region was highly homozygous, with only 17 heterozygous SNPs between the two haplotypes.

The variation in the VSR sizes correlated with differences in intergenic repeat content across *Vitaceae* clades (**Fig. 3**; **Supplementary Table 7**), suggesting that this is the primary factor driving length differences between haplotypes. In the three *Ampelopsis* haplotypes, repetitive elements accounted for 90.2 ± 10.5 kbp, with 19.8 ± 7.1 kbp attributed to LTR *Copia* elements. By contrast, the *Cissus* species exhibited low repeat content, averaging 9.9 ± 3.9 kbp, with notable differences between haplotypes. For instance, haplotypes 1 and 2 of *C. amazonica* and haplotype 4 of *C. gongylodes* contained 6.2 ± 2.2 kbp of LTR *Copia*, while 5.6 kbp of LTR *Gypsy* elements were identified in haplotypes 1, 2, 5, and 6 of *C. gongylodes*. In *P. quinquefolia*, intergenic repeats comprised 50.6 kbp and 45.5 kbp in haplotypes 1 and 2, respectively, with the difference primarily due to the larger amount of LTR *Copia* in haplotype 1 (6.3 kbp) compared to haplotype 2 (1.6 kbp). The greatest amount of intergenic repeats was observed in *T. voinieranum*, with 731.5 and 668.4 kbp in haplotype 1 and 2, respectively. In the outgroup *L. coccinea*, intergenic repeats accounted for 48.4 kbp in both haplotypes.

### The *Vitis* SDR-homologous region shows no evidence of linkage constraint in dioecious *Tetrastigma* species

Given that *Tetrastigma* spp. are dioecious and two VSRs were found in the genome of *T. voinieranum*, we investigated whether the VSR could also be associated with sex determination in *Tetrastigma*. To test this, DNA-seq short reads from thirteen *Tetrastigma* species were aligned to the haplotype 1 of *T. voinieranum* (**Supplementary Table 6**). A total of 3,326,648 SNP positions were identified, of which 15.75% (524,113) were in protein-coding exons. Phylogenetic analysis based on this SNP dataset grouped these species into three clades (**Supplementary Fig. 2**), consistent with previous studies (Peng et al., 2021). The SNP dataset was then used to evaluate the LD in *Tetrastigma* spp. LD declined rapidly, with half of the maximum average r^2^ at 118 bp (**Supplementary Fig. 3**). Concerning the VSR, 3,623 SNP positions were identified across the thirteen species of *Tetrastigma*, with each species averaging 448.1 ± 233.7 SNPs. The ratio between homozygous and heterozygous SNPs was significantly greater in *Tetrastigma* species compared to *M. rotundifolia* and *Vitis* spp. (**Supplementary Fig. 4**; Kruskal-Wallis test followed by a post hoc Dunn’s test; adjusted *P* value < 0.05), indicating lower heterozygosity in *Tetrastigma* spp. in the region. In SDRs, higher heterozygosity is generally expected due to the accumulation of sex-specific alleles, as recombination suppression typically prevents the exchange of genetic material between homologous chromosomes. The observed lower heterozygosity in *Tetrastigma* VSR suggests the absence of sex-specific mutations in the region. To investigate further, we examined the patterns of linkage disequilibrium (LD) in the VSR to assess the presence of any recombination suppression, a common feature in SDRs of dioecious species. No pattern of strong LD could be observed in the VSR, with an average r^2^ of 0.005 ± 0.035 per kbp^2^ (**Fig. 4a**; **Supplementary Fig. 5**). This indicates that the VSR in *T. voinieranum* is not under recombination suppression.

**Fig. 4:**
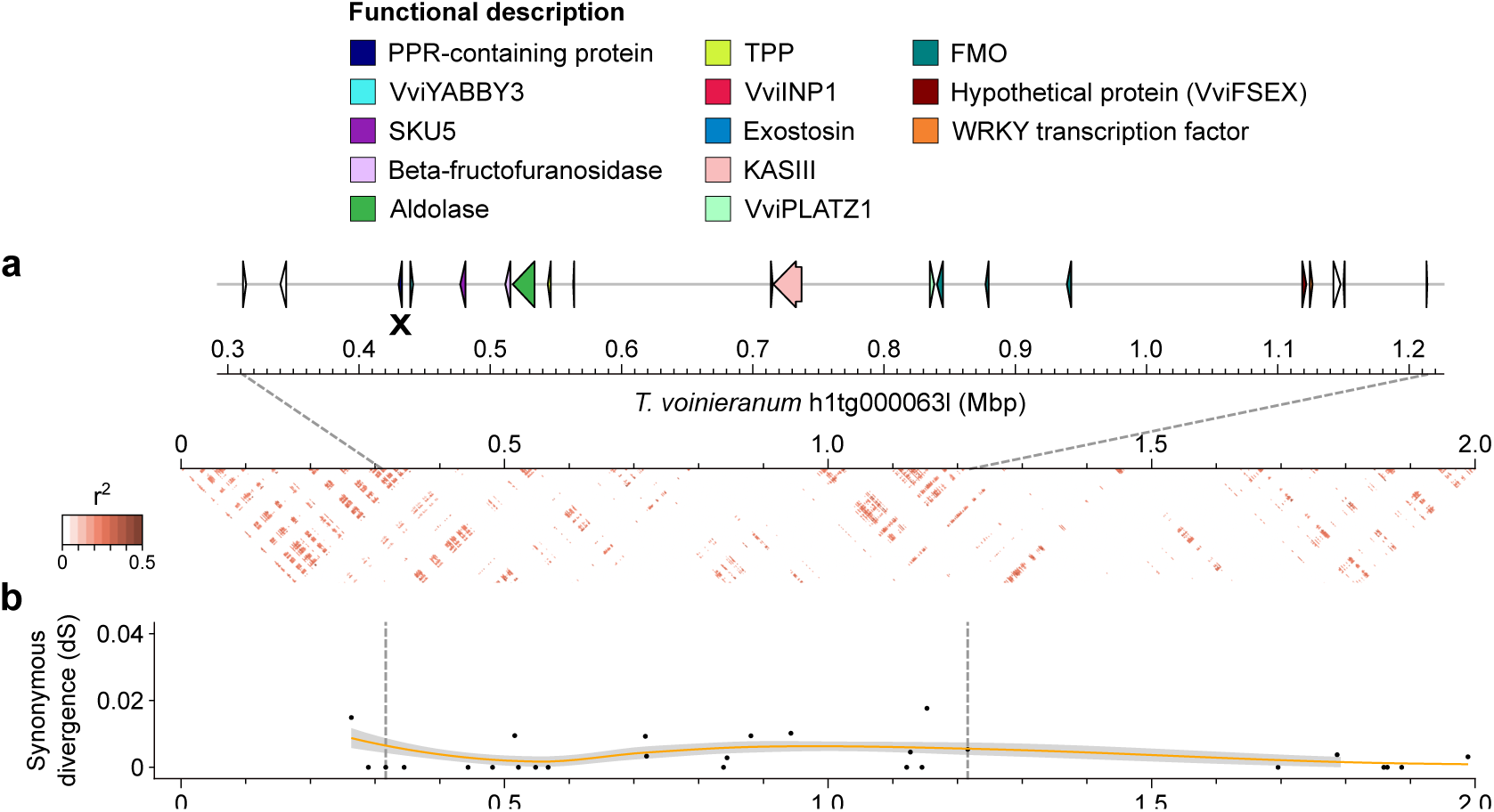
Homologous region of the *Vitis* sex-determining region in *T. voinieranum*. (**a**) Linkage disequilibrium as the mean r^2^ per kbp. Pseudogenes are indicated with an X. (**b**) Synonymous divergence (dS) between the two haplotypes containing the *Vitis*-homologous sex-determining region of *T. voinieranum*.

In addition to evaluating LD, we analyzed sequence divergence between the two VSR haplotypes of *T. voinieranum*. In SDRs, the absence of recombination often leads to increased synonymous divergence (dS) between male and female haplotypes, as genetic material is exchanged less frequently (Charlesworth, 2021). The synonymous divergence (dS) between the two VSR haplotypes of *T. voinieranum* was low and did not show a significant increase in the VSR region (**Fig. 4b**; Kruskal-Wallis test; *P* value = 0.56). This stable dS in the VSR further supports the absence of recombination suppression in this region. Taken together, these results suggest that the VSR in *T. voinieranum* is not associated with sex determination.

### The sex-determining region of *Muscadinia rotundifolia*

To define the boundaries of the muscadine SDR, we first aligned two genetic sex-linked markers (Lewter et al., 2019) on the diploid chromosome-scale genome of *M. rotundifolia* Trayshed (Massonnet et al., 2020). These markers, located at 3,961,704 and 4,493,503 bp on chromosome 2, haplotype 2, identified the F haplotype. Next, we assessed LD in *M. rotundifolia* using whole-genome sequencing data from ten muscadine individuals, three females, three males, and four hermaphrodites (**Supplementary Table 6**). LD decayed rapidly, with half of the maximum average r^2^ observed at 2.3 kbp (**Supplementary Fig. 6**), consistent with findings from previous studies in grapes (Cochetel et al., 2023). Patterns of LD on haplotype 2 of Trayshed’s chromosome 2 revealed elevated LD across the SDR, specifically between 4.268 and 4.414 Mbp, with an average r² of 0.60 ± 0.18 per kbp. This region spans from the vacuolar processing enzyme (VPE) gene to the third *FMO* gene in the F haplotype of the SDR (**Fig. 5a; Supplementary Fig. 7**). In *Vitis* spp., a high LD was observed between *VviYABBY3* and *VviAPT3* (Zou et al., 2021). These results suggest that the boundaries of the *Vitis* and *M. rotundifolia* SDRs are highly comparable.

**Fig. 5:**
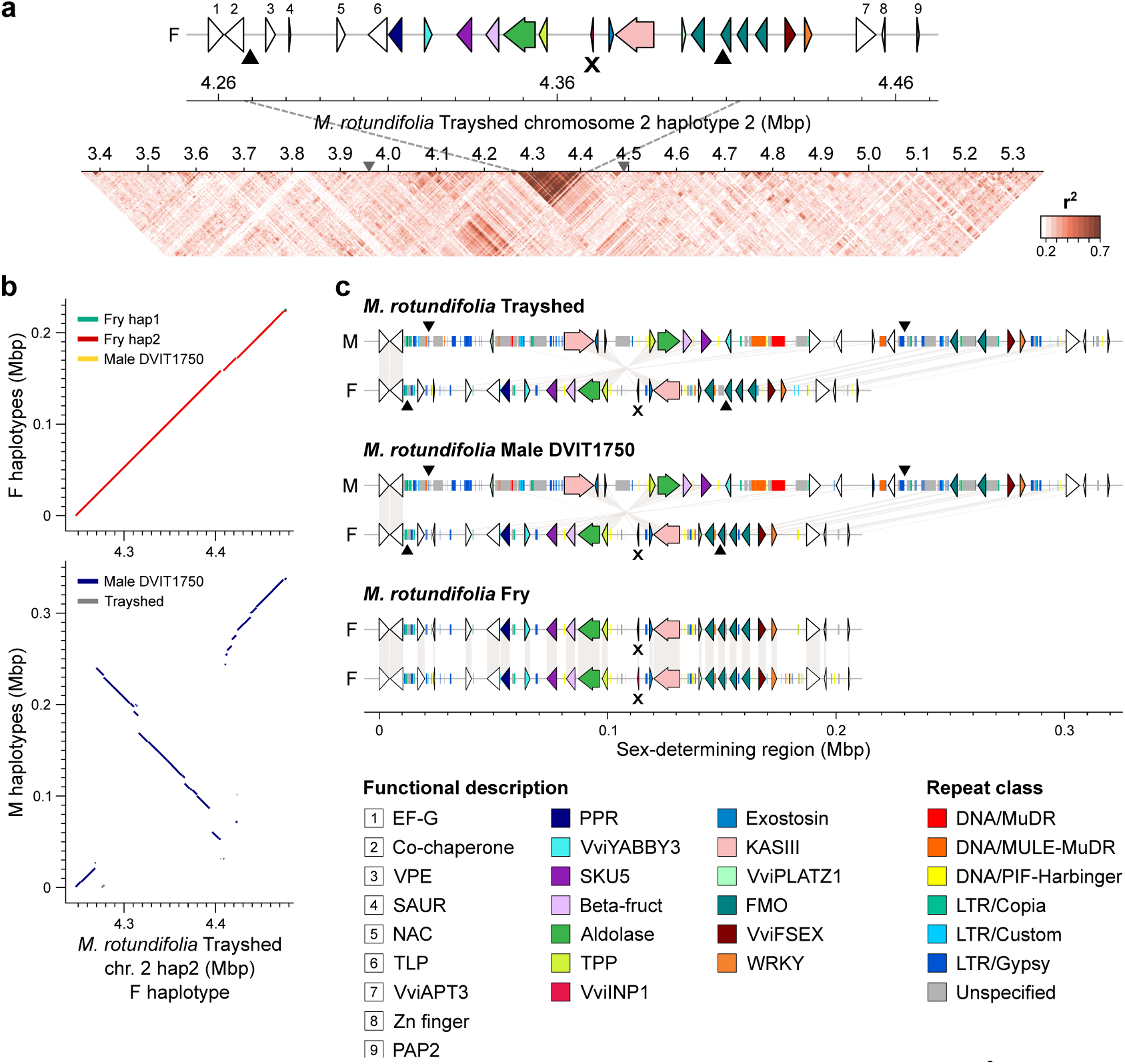
Sex-determining region of *M. rotundifolia*. (**a**) Linkage disequilibrium as the mean r^2^ per kbp. The gray triangles on the scale mark the position of the sex-linked markers in muscadines, S2_4635912 and S2_5085983 (Lewter et al., 2019). (**b**) Whole-sequence alignments of the F (top) and M (bottom) haplotypes against the F haplotype of *M. rotundifolia* Trayshed. (**c**) Schematic representations of the gene and intergenic repeat contents in the sex-determining region (SDR) of the males *M. rotundifolia* Trayshed and DVIT1750, and the female muscadine Fry. For each haplotype, genes are annotated with arrows and repeats with rectangles. White arrows depict genes that are not part of the *Vitis* SDR. The black triangles on the scale mark the position of the breaking points of the inversion in the M haplotype compared to the F haplotype. Pseudogenes are indicated with an X. Gene content representations from panels (**a**) and (**b**) share the same legend.

To investigate the structure of the muscadine SDR, we sequenced and assembled the genomes of the accessions Fry (female) and DVIT1750 (male) using PacBio continuous long reads (CLRs) DNA sequencing (**Supplementary Tables 3-4**). The diploid genome assemblies of Fry and DVIT1750 were 700.9 Mbp and 761.4 Mbp in length, respectively (**Supplementary Table 5**), consistent with the genome size of Trayshed (825 Mbp; (Massonnet et al., 2020). Thanks to the high contiguity of these diploid genomes, we identified the four SDR haplotypes in Fry and DVIT1750. The assignment of these haplotypes to either the F or M haplotype was based on the structure of Trayshed’s haplotypes (Massonnet et al., 2020). Synonymous divergence between the M and F haplotypes of *M. rotundifolia* Trayshed and DVIT1750 was significantly greater at the SDR compared to the adjacent regions (Kruskal-Wallis test; *P* value < 0.05), while no significant difference was observed between the two F haplotypes of Fry (**Supplementary Fig. 8**). These findings support that the muscadine SDR is under recombination suppression.

Alignment of all muscadine haplotypes to Trayshed’s F haplotype revealed that the four F haplotypes were highly similar in structure, as were the two M haplotypes from Trayshed and DVIT1750 (**Fig. 5b**). Both M haplotypes exhibited an inversion of approximately 209 kbp relative to the F haplotype, spanning from 4,506,939 to 4,715,941 bp in the M haplotype of Trayshed. The breakpoints of this inversion closely aligned to the boundaries of strong LD (**Fig. 5a**). Consequently, we extended the region of interest to include two additional genes upstream of the 5’ boundary of the SDR, coding for an elongation factor G (EF-G) and a co-chaperone (**Fig. 5a**). The region spanned 319.8 kbp in the M haplotypes, whereas the F haplotypes averaged 209.6 ± 4.6 kbp. This size difference was largely attributed to the higher content of repetitive elements in the intergenic space of the M haplotypes compared to their F counterparts (**Fig. 5c**; **Supplementary Table 8**).

Repetitive elements accounted for 29.4 ± 3.0 kbp in the F haplotypes and and 125.4 ± 1.1 kbp in the M haplotypes. For example, multiple LTRs were observed in the intergenic region of the M haplotypes, specifically between the co-chaperone gene, *VviPLATZ1* and the gene encoding the 3-ketoacyl-acyl carrier protein synthase III (KAS III), as well as between *VPE* and a *FMO* gene. We estimated the insertion of LTRs between *VviPLATZ1* and *KASIII* in the M haplotypes to have occurred 41.7 ± 10.3 million years ago (Mya) based on their genetic distance. Additionally, several *Mutator*-like elements (MULEs) were found in the intergenic region between *VviYABBY3* and the NAC TF gene in the M haplotypes, which were absent in the F haplotypes. Despite these differences, the gene content in the region was highly conserved between the F and M haplotypes (**Fig. 5b**). The 8-bp deletion in *VviINP1* was present in all F haplotypes and was homozygous in the female Fry, similar to the female *Vitis* individuals.

To estimate when recombination ceased in the grape SDR of *Vitis* and *M. rotundifolia*, we calculated the synonymous divergence (dS) between the M and F allele of each gene within the region for each male individual. The dS values varied considerably along the SDR in both *Vitis* and muscadines (**Supplementary Fig. 9**). In *Vitis* spp., the dS ranged from 0.0034 to 0.0543, while in *M. rotundifolia,* the range was higher, from 0.0174 to 0.064. These results suggest that recombination suppression occurred earlier in *M. rotundifolia* (20.3 ± 8.0 Mya) compared to *Vitis* spp. (14.1 ± 8.2 Mya).

Comparison of the proteins encoded by the genes within the muscadine SDR revealed a sex-linkage pattern from the co-chaperone homologous to *AtHOP1* to the zinc-finger protein (**Supplementary Fig. 10**). When compared with the SDR proteins from *Vitis* spp., twelve out of twenty-one proteins showed a genus-specific separation (**Supplementary Fig. 11**). We observed clustering of the F alleles for *VviINP1* and exostosin genes in both *Vitis* and *M. rotundifolia* relative to the M and H alleles, though no clustering of the M alleles compared to the F and H alleles was observed for any gene.

The two M alleles of *M. rotundifolia* were distinctly separated from all the other alleles for four proteins: VPE, the thylakoid luminal protein (TLP), the aldolase, and the WRKY TF. Next, we compared the promoter region (up to 3kbp upstream of the transcription start site (TSS)) of each SDR gene (**Supplementary Fig. 12**). Similar to the protein level analysis, a sex-linkage pattern was detected for the F alleles of *VviINP1*, exostosin, and *VviPLATZ1* relative to their M and H counterparts. *VviPLATZ1* has been shown to influence grapevine stamen erectness (Iocco-Corena et al., 2021), suggesting that *VviPLATZ1* may play a similar role in *M. rotundifolia*. Notably, the only clear separation between M alleles and the F/H alleles was observed in the promoter region of *VviYABBY3* (**Fig. 6a**; **Supplementary Fig. 12**). To assess the potential impact of the promoter sequence variation of *VviYABBY3* on gene expression regulation, we compared the TF-binding sites. A MYB59-binding site, located 2,104 ± 14 bp upstream of the TSS, was found specific to both *Vitis* and *M. rotundifolia* M alleles, i.e. present in all M haplotypes but absent in all F and H haplotypes. Additionally, a greater number of TF-binding sites in M alleles compared to F and H alleles was identified for seven other TFs (**Supplementary Table 9**), including MYB81 which played a critical role in the developmental progression of microspores in *Arabidopsis thaliana* (Oh et al., 2020), and HOMEODOMAIN GLABROUS 1 (HDG1), involved in floral identity (Kamata *et al*., 2013). In contrast, F and H alleles contained more TF-binding sites for five TFs, such as SPL4 and SPL12, two squamosa promoter-binding protein-like proteins associated with flowering in *A. thaliana* (Jung et al., 2016; Chao et al., 2017). We then quantified the gene expression of the two alleles of *VviYABBY3* in the sexual structures (ovaries and stamens) of the male accessions DVIT1750 and Trayshed, and the female Fry (**Fig. 6b**). In the ovaries, the F allele of *VviYABBY3* was highly expressed (107.2 ± 23.5 transcripts per million (TPM)) in the female flowers of Fry, while its expression was low in the male flowers of DVIT1750 and Trayshed (2.6 ± 0.6 TPM). However, the M allele of *VviYABBY3* was more highly expressed in the ovaries of both male muscadines compared to the F allele (Kruskal-Wallis test; *P* value < 0.05), suggesting a potential role in suppression of femaleness. Regarding the stamens, both alleles were lowly expressed in the three accessions (Fry, 0.2 ± 0.1 TPM; DVIT1750, 1.0 ± 0.8; Trayshed, 1.6 ± 0.2), with the M allele of *VviYABBY3* more highly expressed than the F allele in DVIT1750 (Kruskal-Wallis test; *P* value < 0.05), suggesting a potential role in maleness.

**Fig. 6:**
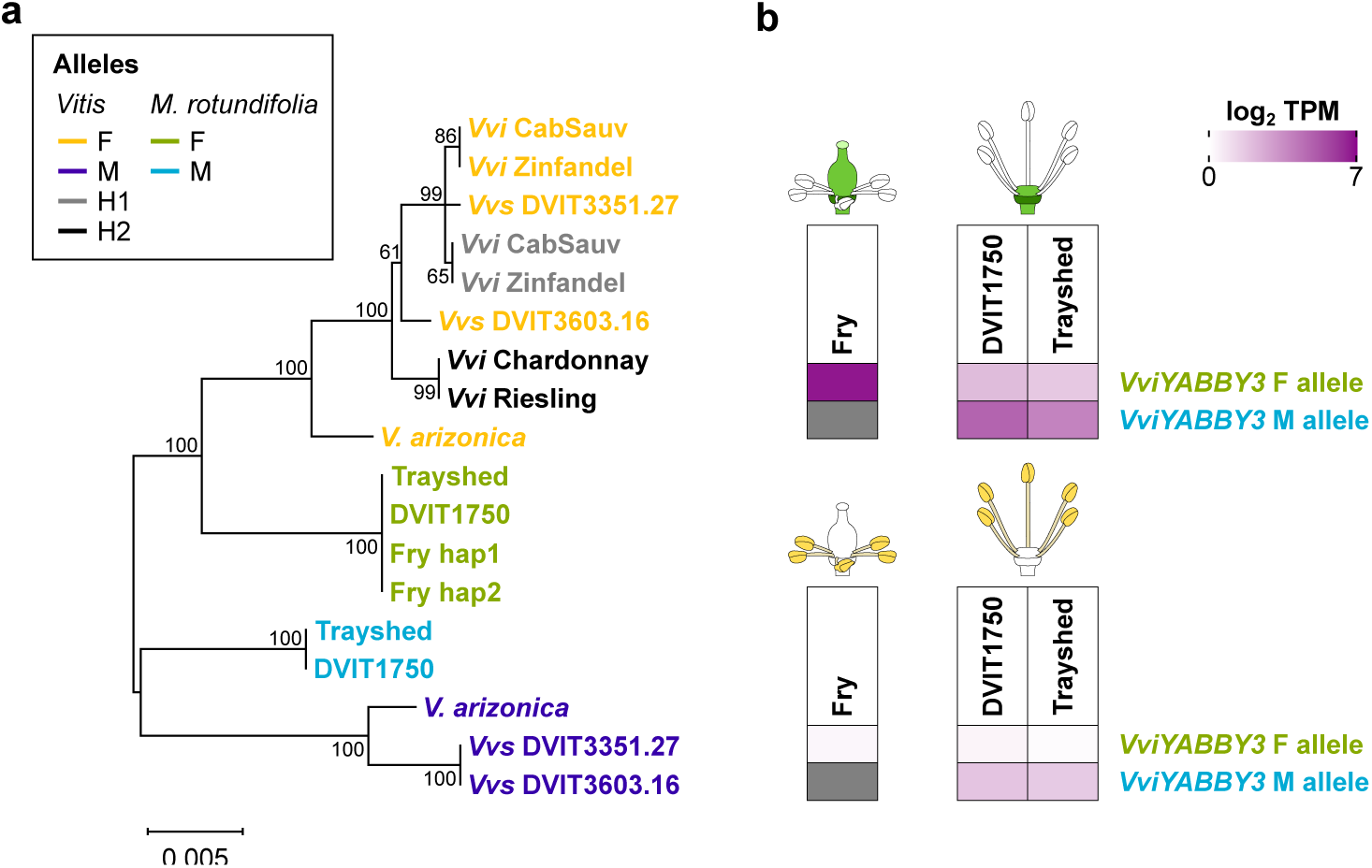
Differences between the F and M allele of *VviYABBY3* in *M. rotundifolia*. (**a**) Phylogenetic tree of the promoter region of *VviYABBY3* in *Vitis* and *M. rotundifolia*. (**b**) Gene expression of the F and M alleles of *VviYABBY3* in flower ovaries and stamens.

## Discussion

### Collinearity and conservation of the *Vitis* SDR in angiosperms

The evolution of sex determination in plants, particularly in dioecious species, is a complex process shaped by genetic changes in regions comprising genes associated with female and male organogenesis (Ming et al., 2007). In garden asparagus (*Asparagus officinalis*), the gene *SUPPRESSOR OF FEMALE FUNCTION* (*SOFF*) is associated with the malformation of the stylar tube and receptive stigma, while *DEFECTIVE IN TAPETAL DEVELOPMENT AND FUNCTION1* (asp*TDF1*) is involved in anther development and pollen production (Harkess et al., 2020). In kiwifruit (*Actinidia deliciosa*), the female-suppressing gene, *Shy Girl* (*SyGI*), represses the development of the pistil through negative regulation of the cytokinin signaling, and the fasciclin-like gene *Friendly Boy* (*FrBy*) is involved in proper tapetum degradation (Akagi et al., 2019). Concerning the *Vitis* SDR, at least five genes within the locus, including the two candidate sex-determining genes *VviYABBY3* and *VviINP1*, are, or have homologous genes, involved in floral organ development and identity in plants. Mutations in homologous genes of the *Vitis* SDR locus in Arabidopsis have been shown to cause significant alterations in flower morphology and fertility (**Table 1**), highlighting the functional importance of these genes in proper floral development.

The high conservation and collinearity of the genes within the *Vitis* SDR locus among angiosperms suggest that this region has been under strong evolutionary pressure to maintain its structure and function (**Fig. 2**) (Tang et al., 2008). The higher rank-normalized gene conservation scores further imply that not only the arrangement but also the sequence of these genes is highly preserved, reinforcing the idea that this locus plays a critical role in flowering processes (Nair et al., 2022). The absence of the *Vitis* SDR in non-flowering plants further supports its specialized role in angiosperm reproductive development (Pellegrini et al., 1999). The *Vitis* SDR is the only plant SDR shown to have high gene collinearity and conservation across angiosperms to date. Similarly, the male-determining gene *FrBy* from kiwifruit has been shown to have orthologs in 32 angiosperm species, including three with similar gene functions, suggesting that the function of *FrBy* is conserved across flowering plants (Akagi et al., 2019). The roles of other SDRs in flowering processes, beyond defining sex dimorphism, remain an open area for exploration.

### Gene content of *Vitis* SDR-homologous regions is highly conserved in Vitaceae

Among Vitaceae, we observed variation in haplotype size mainly due to differences in the repeat content of intergenic regions, with types of repetitive elements varying between species. Although most of the genes within the locus were conserved, some differences in gene content were evident at the clade, species, and individual levels. In the outgroup *L. coccinea*, two VSR haplotypes were identified, both similar in size and gene content, with only a few SNPs between them, indicating that the region is highly homozygous. Across Vitaceae, the size of the VSRs varied significantly between clades, ranging from 118.2 kbp in *C. amazonica* to 902.6 kbp in *T. voinieranum* (**Fig. 3**).

### Emergence of dioecy in Vitaceae: flower sex is determined by different mechanisms in grapes and *Tetrastigma* spp

Because the genus *Tetrastigma* from the Cayratieae clade is dioecious, and *T. voinieranum* genome contained two VSRs, we investigated if the region could also be associated with sex determination. A requisite for the SDR formation is the linkage constraint between the two sex-determining genes (Charlesworth, 2013). Evaluation of the LD in the region using short DNA reads from thirteen *Tetrastigma* species showed a low LD, and no pattern of recombination suppression could be observed. Low LD indicates that the region is unlikely associated with sex determination, suggesting that dioecy emerged in two distinct regions in the Cayratieae and Viteae clades. This supports the hypothesis that dioecy arose independently in the Vitaceae family based on their geographical locations (Gerrath et al., 2015). While *Vitis* spp. is found in the temperate northern hemisphere, *Tetrastigma* spp. is located in tropical southern hemisphere areas. Sequencing additional *Tetrastigma* species and varieties would help to validate the absence of linkage constraint in the region. Coupled with a flower sex type phenotyping, this might allow to identify the *Tetrastigma* SDR.

### Dioecy emerged before the divergence of *Vitis* and *M. rotundifolia* genera

From evolutionary studies, the divergence between *Vitis* and *Muscadinia* genera was estimated between 18 and 47 Mya (Wan et al., 2013; Liu et al., 2016; Ma et al., 2018). In this study, we aimed to better characterize the muscadine SDR to assess the commonalities and differences between the two grape SDRs. The identification of the muscadine SDR using sex-associated genetic markers and pattern of LD showed that the SDR of *M. rotundifolia* was located in a region similar to the *Vitis* SDR. This suggests that the dioecy emerged before the divergence of the two grape genera. Additional sequencing data from *M. rotundifolia*, such as recombinants, will be necessary to confirm and narrow down the muscadine SDR boundaries. This same approach was successful in determining the boundaries of the *Vitis* SDR (Zou et al., 2021). As a result of breeding efforts, two independent hermaphroditic lines, designated H1 and H2, have been identified in muscadines (Dearing, 1917). Comparing the SDR from additional male, female, and hermaphrodite *M. rotundifolia* would thus help to refine the boundaries of the region and better understand how hermaphroditism arose. A large inversion was found in the M haplotype of both male muscadines relative to the F counterpart. In *Carica papaya*, two large inversions were also observed in the male- and hermaphrodite-specific region of the Y chromosome compared to the X region (Wang et al., 2012a; VanBuren et al., 2015). Both inversions were found to have caused recombination suppression in the papaya SDR, suggesting that the inversion in the M haplotype of *M. rotundifolia* play<u>s</u> a similar role in the muscadine SDR.

Regarding the candidate sex-determining genes of *M. rotundifolia*, some evidence suggest that they could be the same genes that the ones previously discovered in *Vitis* spp. (Massonnet et al., 2020; Zou et al., 2021). Like in *Vitis* spp., a homozygous 8-bp deletion was found in *VviINP1* in the female *M. rotundifolia* Fry. Pollen grains of female flowers from muscadines, including the accession Fry, were reported as acolporated and sterile (Biasi and Conner, 2016). This suggests that mutations in *VviINP1* could also be the female-determining factor in muscadines. Regarding the female-suppressing factor, the promoter of *VviYABBY3* was the only sequence showing a M linkage in both *M. rotundifolia* and *Vitis* genera (**Fig. 6a**). Comparison of the TF-binding sites between M and F alleles pointed several sites specific to the M alleles. In addition, the M allele of *VviYABBY3* was detected as more highly expressed compared to its F counterpart in ovaries (**Fig. 6b**), suggesting that this allele could also be the female-suppressing gene in *M. rotundifolia*. Finally, visualization of the alignment of DNA-seq short reads from 159 *Vitaceae* accessions, showed that the mutation was absent in all accessions, including the two other Viteae, *Ampelocissus* and *Pterisanthes* (**Supplementary Table 7**). This suggests that the candidate male-sterility mutation is specific to the *Vitis* and *Muscadinia* genera. However, DNA sequencing of additional samples from the hermaphrodite Vitaea genera *Ampelocissus*, *Pterisanthes*, and *Nothocissus*, will be necessary to confirm it.

### Model for the evolution of dioecy in *Vitis* and *Muscadinia*

Our results support a model where, in an ancestor of *Vitis* and *Muscadinia*, mutations within a highly conserved region associated with flowering led to the emergence of dioecy in this lineage (**Fig. 7**). To our knowledge, this is the first observation of sexual dimorphism emerging through the evolution of a genomic region which gene content is highly collinear and conserved in angiosperms and composed of multiple genes playing a role in flower development and morphology, and sexual fertility.

**Fig. 7:**
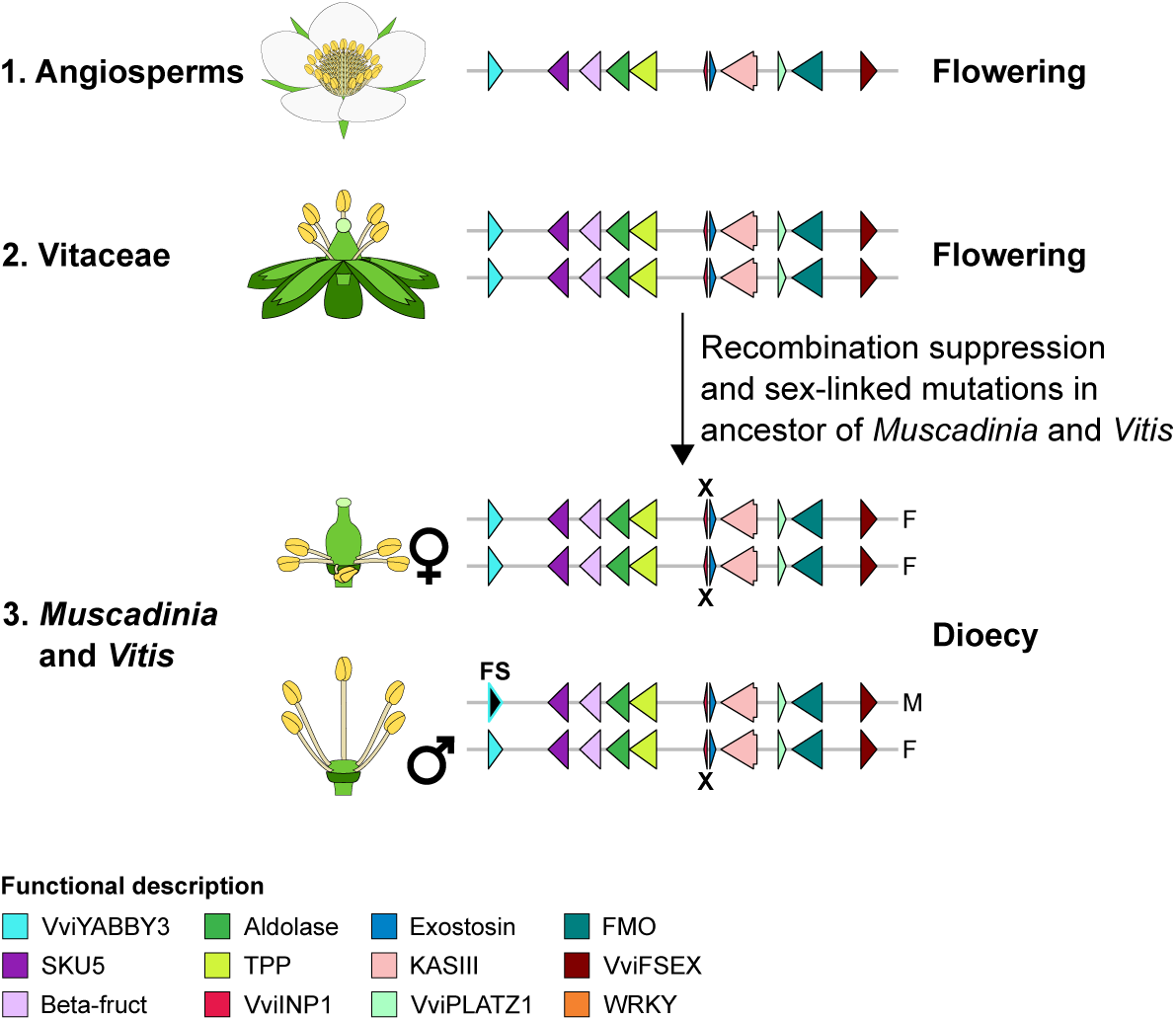
Model of the establishment of the *Vitis* and *Muscadinia* sex-determining regions in a highly conserved and flowering-associated region in angiosperms. Schematic representations of the gene content in the homologous regions of the *Vitis* sex-determining region in angiosperms (1), *Vitaceae* (2), and *Muscadinia* and *Vitis* genera (3). Arrows depict genes. The nonsense mutation in the candidate male-determining in grapes, *VviINP1*, is indicated with an X. The candidate female-suppressing (FS) gene *VviYABBY3* is also indicated.

## Materials and Methods

### Plant material

For genome sequencing, young leaves were collected from 16 accessions: two *M. rotundifolia* individuals, Fry and DVIT1750, six *Ampelopsis* individuals representing four species, six *Cissus* species, one *P. quinquefolia* DVIT2400, *T. voinieranum*, and *L. coccinea* 1464. Information about the sequenced accessions is provided in **Supplementary Table 3**. For RNA-sequencing, inflorescences from *M. rotundifolia* Fry (female), *M. rotundifolia* Trayshed (male), and *M. rotundifolia* DVIT1750 (male), were collected at full flowering with 50% caps off (E-L 23 (Coombe, 1995)). Ovaries and stamens from cap-off flowers were then sampled to represent three biological replicates per accession. All plant material was immediately frozen after collection and ground to powder in liquid nitrogen.

### DNA and RNA extraction, library preparation, and sequencing

For long-read DNA sequencing, high molecular weight genomic DNA (gDNA) was isolated as in Chin *et al*. (2016) from *M. rotundifolia* Fry and DVIT1750, *A. aconitifolia* DVIT2492, *A. glandulosa* var. brevipedunculata PI 597579, *C. amazonica*, *C. gongylodes*, *P. quinquefolia* DVIT2400, *T. voinieranum*, and *L. coccinea* 1464. DNA purity and quantity were evaluated using a Nanodrop 2000 spectrophotometer (Thermo Scientific, IL, USA) and a Qubit™ 1X dsDNA HS Assay Kit (Thermo Fisher, MA, USA). DNA integrity was assessed with the DNA High Sensitivity kit (Life Technologies, CA, USA) and by pulsed-field gel electrophoresis. SMRTbell libraries of *M. rotundifolia* Fry and DVIT1750, *A. aconitifolia* DVIT2492, *A. glandulosa* var. brevipedunculata PI 597579, *C. amazonica*, *C. gongylodes*, and the HiFi libraries of *P. quinquefolia* DVIT2400, *T. voinieranum*, and *L. coccinea* 1464 were prepared as described in Minio *et al*. (2019; 2022), and sequenced using PacBio Sequel II system (Pacific Biosciences, CA, USA) at the DNA Technology Core Facility, University of California, Davis (Davis, CA, USA). Summary statistics of long-read DNA sequencing are provided in **Supplementary Table 4**. For short-read DNA sequencing, DNA extraction and library preparation were performed as Massonnet *et al*. (2020). Regarding the RNA-sequencing from the muscadine flowers, RNA extraction and library preparation were performed as in Rapicavoli *et al*. (2018). RNA extraction for one bioreplicate of stamens from Trayshed flowers failed and tissue from the bioreplicate was not sufficient to repeat the extraction. DNA and cDNA libraries were sequenced using an Illumina HiSeqX Ten system (IDseq, Davis, CA, USA) as 150-bp paired-end reads and an Element Bio AVITI as 80-bp paired-end reads, respectively.

### Genome assembly and annotation

Genome assembly from PacBio CLR reads was performed with FALCON-Unzip (Chin et al., 2016) as Massonnet *et al*. (2020) and with hifiasm (Cheng et al., 2021) as Minio *et al*. (2022a) for the HiFi reads. The repeat and gene annotations were made as described in Massonnet *et al*. (2020). Gene space completeness was evaluated using BUSCO v.5.4.7 and the library embryophyte_odb10 (Waterhouse et al., 2018). For the genome assemblies of the *Ampelopsis*, *Cissus*, *Parthenocissus*, *Tetrastigma*, and *Leea* species, repeat libraries were created separately with RepeatMasker v.open-4.0.6 (Smit et al., 2013) and RepeatModeler2 v.2.04 (Flynn et al., 2020) starting from the RepBase library v.20160829 and Cabernet Sauvignon custom repeat library (Minio et al., 2019b). For the gene annotation, we used as experimental evidence the protein-coding sequences and the proteins of the isoforms identified using IsoSeq data from Cabernet Sauvignon (Minio et al., 2019b), North American *Vitis* species (Cochetel et al., 2023), and *M. rotundifolia* Trayshed (Cochetel et al., 2021), as well as the transcriptome of *Vitis vinifera* ssp. *vinifera* Corvina (Minio et al., 2019b). Sequence redundancy was reduced using CD-HIT v4.6 (Li and Godzik, 2006) with the parameters “cd-hit-est -c 0.95”.

### Phylogenetic analysis

Phylogenetic tree of the eleven *Vitaceae* and *L. coccinea* (**Fig. 1**) was based on the 353 putatively single-copy protein-coding genes in angiosperms (Johnson et al., 2019). Orthologues of the proteins encoded by the *Arabidopsis thaliana* genes from the Angiosperms353 set (Johnson et al., 2019) were identified among the twelve genomes. Genomes of *V. vinifera* ssp. sylvestris O34-16 (female) and DVIT3351.27 (male), and *M. rotundifolia* Trayshed were retrieved from Massonnet et al. (Massonnet et al., 2020) and Massonnet et al. (Massonnet et al., 2022), respectively. Proteins of *A. thaliana* were aligned and annotated on the primary and haplotype 1 contigs using miniprot v.0.4-r174-dirty (Li, 2023). For each protein, the alignment with the greatest coverage and identity, with no frameshift nor stop codon, was retained. Protein sequences were generated using gffread from Cufflinks v.2.2.1 (Trapnell et al., 2010). Multi-sequence alignments of the proteins were performed with MUSCLE v.3.8.31 (Edgar, 2004). Alignments were then concatenated and parsed using Gblocks v.91b (Castresana, 2000) with up to 8 contiguous non-conserved positions, a minimum length of a block of 10, and no gap. Phylogenetic analysis was conducted with MEGAX (Kumar et al., 2018) using the Maximum Likelihood method, the JTT matrix-based substitution model (Jones et al., 1992) with four Gamma categories, and 1,000 replicates. For inferring the divergence times between genera, the RelTime method (Tamura et al., 2018) was applied using three calibration points corresponding to the divergence time between (i) the *Vitis* and *Muscadinia* spp. (41.58 million years ago), (ii) the Viteae and Parthenocisseae clades (59 million years ago), (iii) the *Tetrastigma* and *Cissus* spp. (72.75 million years ago) (Herrera et al., 2024), with a normal distribution and a standard deviation of 1.

Other phylogenetic analyses were performed with MEGAX (Kumar et al., 2018) using the Neighbor-Joining method (Saitou and Nei, 1987) and 1,000 replicates. The Poisson correction method (Zuckerkandl and Pauling, 1965) and the Kimura 2-parameter method (Kimura, 1980) were used for generating the trees based on protein and DNA sequence comparison, respectively. Prior the phylogenetic tree of the thirteen *Tetrastigma* species, a sequence alignment matrix was created from the SNPs identified between the thirteen species and the haplotype 1 of *T. voinieranum* genome using the script vcf2phylip.py v.2.0 (Ortiz, 2019).

### Gene collinearity analysis

Gene collinearity was evaluated between the haplotype 1 of Cabernet Sauvignon genome (Massonnet et al., 2020) and 42 plant genomes. Information about the genome and annotation files used for this analysis can be retrieved in **Supplementary Table 1**. Proteins of each plant accession were aligned in a pairwise fashion against each other using DIAMOND blastp v2.0.13.151 (Buchfink et al., 2021) with the following parameters: “--max-target-seqs 100 --evalue 0.001”. Colinear gene loci were identified with MCScanX v.11.Nov.2013 (Wang et al., 2012b) and the parameters “-s5 -m25 -w1”. Gene content of the haplotype 1 of Cabernet Sauvignon was then separated in window of 20 consecutive genes. For each plant genome, the collinear blocks with the highest number of genes from the Cabernet Sauvignon 20-gene windows were selected to form collinear windows. For this analysis, the grape SDR was considered as the region spanning from the NAC TF gene to the gene homologous to *A. thaliana* PURPLE ACID PHOSPHATASE 2 (*PAP2*). A gene conservation score was assessed for each collinear gene in angiosperms as described in Nair *et al*. (Nair et al., 2022). Gene conservation score was calculated by dividing the alignment score (bit score) between collinear genes by the bit score of the grape reference protein against itself, resulting in values between 0 and 1. Gene conservation scores were then rank-normalized across superrosids, superasterids, and monocots, separately, to control for the evolutionary distance between the reference and each angiosperm clade (**Supplementary Fig. 13**). Genes absent from the syntenic window corresponding to the *Vitis* SDR were searched using miniprot v.0.4-r174-dirty (Li, 2023) and the SDR protein sequences of the haplotype 1 of Cabernet Sauvignon.

### Sex-determining region localization and haplotype reconstruction

Homologous regions of the grape SDR were identified by aligning the protein-coding sequences (CDS) from the SDR of Cabernet Sauvignon Hap1 onto the nine genome assemblies with GMAP v.2020-06-01 (Wu and Watanabe, 2005). For *M. rotundifolia*, the closest markers to the SDR, S2_4635912 and S2_5085983 (Lewter et al., 2019), were identified as described in Cochetel *et al*. (Cochetel et al., 2021). When the alignments of the SDR-associated sequences were found on multiple contigs, NUCmer from MUMmer v.4.0.0 (Marçais et al., 2018) and the “--mum” option were used to determine the overlap between contigs. Then, contigs were reconstructed using HaploMake from the tool suite HaploSync v.1.0 (Minio et al., 2022b). Gene models among the grape SDR-like regions were manually refined using the intron-exon structure of the SDR-associated genes from Cabernet Sauvignon. Schematic representations of the gene content were made using the R package gggenes v.0.4.1 (https://wilkox.org/gggenes/).

### Alignment of short DNA sequencing reads

Because of sequencing depth variability, samples from the muscadines and the *Tetrastigma* species were randomly subsampled to 50 and 30 million reads, respectively, with seqtk sample from the package seqtk v.1.2-r101-dirty (https://github.com/lh3/seqtk) and the parameter -s100 before trimming. Short DNA-seq reads were trimmed using Trimmomatic v.0.36 (Bolger et al., 2014) and the following settings: “LEADING:3 TRAILING:3 SLIDINGWINDOW:10:20 MINLEN:33”. High-quality paired-end reads were aligned onto their corresponding reference genome using bwa v.0.7.17-r1188 (Li and Durbin, 2009) (**Supplementary Table 6**). Alignments were visualized using Integrative Genomics Viewer v.2.4.14 (Robinson et al., 2011) to evaluate the zygosity status of the 8-bp deletion in *VviINP1*.

### Variant calling

Prior variant calling, PCR and optical duplicates were removed with Picard tools v.2.8 (http://broadinstitute.github.io/picard/). The variant calling was performed using HaplotypeCaller from GATK v.4.2.2.0 (DePristo et al., 2011) with the parameters “--sample-ploidy 2 -ERC GVCF”. VCF files were combined and genotyped using the programs CombineGVCFs and GenotypeGVCFs from GATK v.4.2.2.0 (DePristo et al., 2011) with default parameters. SNPs with a quality higher than 30, a depth of coverage higher than five reads, no more than three times the median coverage depth across accessions, a minor allele frequency higher than 0.05, and no missing data among individuals were filtered with vcftools v0.1.15 (Danecek et al., 2011). For *Tetrastigma* genus, SNPs were filtered using a depth of coverage higher than one read and lower than 15 reads due to the low sequencing coverage (3.1 ± 0.6 X). SNPs were further filtered using the “filter” function of bcftools v.1.9 (Danecek et al., 2021) and the options “-e QD < 2.0 | FS > 60.0 |MQ < 40.0 | MQRankSum < −12.5 | ReadPosRankSum < −8.0 | SOR > 3.0”. SNPs within the muscadine SDR were used to perform a principal components analysis using Plink v.1.90b5.2 (Purcell et al., 2007) to infer the flower sex type of the samples SRR6729328, SRR7819188, SRR7819190, SRR11886267 (**Supplementary Fig. 14**).

### Linkage disequilibrium analysis

To assess linkage disequilibrium (LD) decay, LD was estimated using Plink v.1.90b5.2 (Purcell et al., 2007) and the following command: “plink --vcf name.vcf --double-id --allow-extra-chr -r2 gz --maf 0.05 --ld-window 10 --ld-window-kb 300 --ld-window-r2 0 --out name”. SNPs from the muscadines were randomly subsampled using the parameter “--thin-count 800000”. LD decay was evaluated using the model from Hill and Weir (Hill and Weir, 1988). To explore the LD landscape of the SDR, we used Tomahawk v.beta-0.7.1 (https://github.com/mklarqvist/tomahawk). SNPs comprised in the 2-Mbp region around the SDR in muscadines and *Tetrastigma* spp. were used as input to calculate the LD with the “calc” function. The r^2^ values were aggregated into bins of 1 kbp using the “aggregate” function, the parameters “-r mean -c 5”, and with 2,000.

### Whole-sequence alignments and structural variation analysis

Pairwise sequence alignments were performed using NUCmer from MUMmer v.4.0.0 (Marçais et al., 2018) and the “--mum” option.

### Divergence time estimation

For each male *Vitis* and *M. rotundifolia* individual, the coding sequences of the M and F alleles of each SDR gene were aligned using MUSCLE v.3.8.31 (Edgar, 2004). Synonymous divergence (dS) was calculated using the yn00 program in the PAML package v.4.9 (Yang, 2007). The heatmap of the dS values was generated using geom_tile from the R package Tidyverse 2.0.0 (Wickham et al., 2019). LTR retrotransposons were detected using LTRharvest from the GenomeTools package v.1.6.5 (Gremme et al., 2013). The two LTRs of each LTR retrotransposon were aligned with MEGAX (Kumar et al., 2018) and their genetic distance was estimated using Kimura’s two-parameter model (Kimura, 1980). Divergence time was calculated as T = K/2μ × generation time, where K is the genetic distance and μ is the mutation rate. A generation time of 3 years and a nucleotide substitution rate of 2.5 × 10^−9^ substitutions per base per year were assumed.

### Transcription factor-binding site analysis

For each haplotype, promoter sequences were extracted with a maximum of 3 kbp upstream regions from the gene transcriptional start sites. TF-binding sites were identified using the R packages TFBSTools v.1.38 (Tan and Lenhard, 2016) and the JASPAR2018 v.1.1.1 (Tan, 2017).

### Gene expression analysis

Samples were randomly subsampled to 25 million reads with seqtk sample from the package seqtk v.1.2-r101-dirty (https://github.com/lh3/seqtk) and the parameter -s100 before trimming. RNA-seq reads were trimmed using Trimmomatic v.0.36 (Bolger et al., 2014) and the following settings: “LEADING:3 TRAILING:3 SLIDINGWINDOW:10:20 MINLEN:36”. Transcript abundance was assessed using Salmon v.1.5.1 (Patro *et al*., 2017) and these parameters: “--gcBias –seqBias --validateMappings”. For each accession, a transcriptome index file was built using Trayshed protein-coding sequences ver2.2 and the Trayshed genome v.2.1 (Massonnet et al., 2022) as decoy, and a k-mer size of 31. Counts were imported using the R package tximport v.1.20.0 (Soneson et al., 2016) and combined at gene-level for genes with alternative transcripts. Statistics about the RNA-seq analysis can be retrieved in **Supplementary Table 10**. Because of a low percentage of reads aligning on Trayshed protein-coding sequences, the third bioreplicate of ovaries from DVIT1750 was removed from the dataset. Gene expression of the two alleles of *VviYABBY3* represented in **Fig. 6b** corresponds to the average of the log_2_-transformed (TPM + 1) detected in the three bioreplicates.

## Data availability

Sequencing data and genome sequences are accessible through NCBI under the BioProject PRJNA1151724. Gene annotation files are available at Zenodo doi:10.5281/zenodo.13362874.

## Acknowledgments

We would like to thank Bernard Prins and Claire Heinitz (National Clonal Germplasm, USDA-ARS) for providing information about the *Vitaceae* material, Ernesto Sandoval (University of California Davis) and Daniel Pfarr (Sacramento State University) for providing leaf material from *L. coccinea* 1464, Malin Petersen (University of British Columbia, Canada) for her help during the plant tissues collection.

## Author contributions

M.M. and D.C. designed the project. M.M. and J.L. collected the plant material. R.F.-B. extracted DNA and RNA, and prepared sequencing libraries. M.M and A.M. assembled the genomes. M.M., N.C, A.M., and V.R. performed the data analyses. M.M. and D.C. wrote the manuscript.

